# Mycobacterial load and direct contact primes spacious to compact phagosome switching for egress

**DOI:** 10.1101/2024.09.04.611260

**Authors:** Aher Jayesh Bhausaheb, Aniruddha Nagarajan, Bommidi Jahnavi, Debraj Koiri, Jafarulla Shaikh, Sandeep Choubey, Varadharajan Sundaramurthy, Mohammed Saleem

## Abstract

*Mycobacterium tuberculosis* (*Mtb*) establishes intracellular niches by remodeling host membranes into either spacious or compact phagosomes, yet how direct bacterial contact within these distinct compartments facilitates bacterial egress remains unknown. Using fixed cell imaging, in vitro reconstitution of phagosome-like vesicles, and numerical simulations, we uncover that mycobacterial load and its direct contact determines phagosome fate by driving membrane bending, lipid wrapping and phase separation. Low-to-moderate load induces membrane vesiculation, generating compact phagosome-like structures. High bacterial load drives a scaffold-like architecture that primes compartments for rupture via complete lipid phase separation and changes in the membrane’s bending and area stretch moduli, rendering it more deformable. Notably, physical contact synergizes with the virulence factor ESAT-6, amplifying its membrane-deforming activity to disrupt phagosomal integrity. We propose that mycobacteria actively switch spacious to compact phagosomal states by modulating host membrane mechanics—a process governed by the membrane-to-contact area ratio and proximity to lipid demixing point. These findings reveal a biophysical switch in pathogen-driven membrane remodeling, with broad implications for understanding how intracellular pathogens manipulate host membranes to survive.

## Introduction

The interaction of bacteria with the cellular membrane interfaces is essential for its host entry, intracellular life as well as egress ^1, 2^.Such highly complex processes are driven by multiple coordinated protein or lipid-dependent noncovalent interactions^3^. Several invasive bacterial pathogens, such as *Listeria monocytogenes*, *Pseudomonas aeruginosa* are known to hijack host membrane interfaces for actin polymerization resulting in the bacterial engulfment by the host cell membrane^4, 5^. *P. aeruginosa* induces cell membrane invaginations via a lipid zipper formation by the interaction between the glycosphingolipid Gb3 and the bacterial surface lectin LecA, in addition to other cell surface receptors^6^. Clathrin and Arp2/3 complexes have been identified as the key regulator of actin polymerization for zippering pathogens such as *Listeria* and *Yersinia*^7, 8^. Likewise, *Mycobacterium tuberculosis* (*Mtb*) is known to gain entry into the host cell via passive internalization or engulfment mediated by several surface adhesion proteins^9, 10^. The *Mtb* are ejected from the host cell by generating actin-based structures called ejectosomes that are aided by autophagy machinery enabling a non-lytic mechanism ^11, 12^. Our understanding of bacterial host entry has been reasonably refined in recent times and mediated by a diverse set of pathogen-specific biochemical interactions. However, the disruption of the pathogen-containing vacuolar compartment of the host cell remains an open question, particularly in the case of *Mtb*^13^.

*Mtb* survives intracellularly by hijacking the phagosome maturation ^14, 15^. ESAT-6 (6kDa early secretory antigen target), a highly immunogenic effector protein secreted by *Mtb*, is critical for the process of phagosome (also referred to as Mycobacteria containing vacuole (MCV)) rupture ^16, 17^. Purified recombinant ESAT-6 is known to induce lysis of cultured macrophages through permeabilization of both the phagosome membrane as well as cell membrane upon gaining access to the cytoplasm ^18–20^. Despite significant evidence that supports the direct role of ESAT-6 in phagosome lysis and virulence^21–23^, a recent study reported that ESAT-6 interaction alone is not sufficient for phagosome disruption. Instead, direct bacterial physical contact is important for gross phagosomal membrane disruption ^24^. *In vivo* observations also suggest that ESAT-6 required additional factors such as the lipid phthiocerol dimycocerosates (PDIMs), which are complex branched cell-wall-associated lipids known to induce sterol clustering in host membranes, eventually inducing phagosome disruption ^25–30^. Interestingly, *Mycobacterium marinum* transposon mutants induce phagosome damage despite their inability to secrete ESAT-6, suggesting that ESAT-6 alone may not be sufficient for the ESX-1 virulence function ^31^. Though the ability to damage the phagosomal membrane has emerged central to *Mtb* virulence, the mechanism of phagosome membrane remodelling and disruption remains poorly understood ^13^. *Mtb* is known to be a non-motile bacteria, however, exhibits sliding mobility upon encountering surfaces. Very recently, motile swimming bacteria such as *E. coli* and *B. subtilis* have been shown to induce non-equilibrium membrane fluctuations ^32^, deformations ^33^ as well as conferring coupled mobility in synthetic giant vesicles encapsulating the bacteria^34^. However, the dynamics and impact of direct physical contact of a non-motile bacteria with the vacuolar compartment remain unknown to date. Understanding the implications of direct physical contact of the bacteria is not only important for vacuolar disruption but also for a broad class of bacterial pathogens interaction with host cell membranes both during internalization as well as egress.

In this work, we developed an *in vitro* reconstitution method to dissect the mechanism of phagosome membrane remodelling induced by the direct physical contact of bacteria by encapsulating live *Mycobacterium smegmatis* (as a model opportunistic bacteria) inside a host phagosome-like compartment. Furthermore, it also enables us to delineate the role of direct physical contact in phagosome membrane remodelling from virulence factors. We capture the dynamics of bacterial direct contact as well as bacterial-load-mediated host membrane remodelling. We discover that the direct-contact of an isolated *M . smegmatis* with the encapsulating host membrane results in significant lipid wrapping, membrane bending, and local lipid phase separation at the contact site. In contrast, the spacious and compact phagosome mimic with varying bacterial load induces complete lipid phase separation of the compartment. The lipid wrapping is driven by the surface condensation of the compartmental lipids onto the bacterial surface in direct contact and lowers host lipid diffusion by half at the contact area. The direct physical contact under moderate load decreases the area stretch and bending modulus of the host membrane. On the contrary higher bacterial load induces a scaffold-like architecture priming the host compartment for rupture. The degree of bending and area expansion of the phagosome-like compartment is dictated by the ratio of the compartmental surface area available to the surface area of the bacterial contact site. Further direct contact of the bacteria enhances the membranolytic potential of ESAT-6 thus contributing to its virulence. Together we propose that load mediated direct contact of the bacteria drives the switching from spacious to compact phagosome.

## RESULTS

### Internalization of *Mycobacterium tuberculosis* leads to heterogenous compartmentalization

To capture the dynamics of progression of phagosome maturation during mycobacterial infection, we monitored lysosomal marker acquisition in THP1 macrophages infected with live mCherry-expressing *Mycobacterium tuberculosis* (*Mtb*) for different time periods ranging from 2 to 48 hours post-infection (Fig.1a). Over the course of infection, the proportion of compact phago-lysosomes decreased by nearly 50%, while the number of spacious phagosomes stayed relatively constant (Fig. 1b). Interestingly, between 2 and 48 hours post-infection, an increase in compact phago-lysosome coincided with a decrease in spacious phago-lysosome, revealing an inverse relationship in their distribution (Fig. 1d, e). This dynamic redistribution suggests that spacious phagosomes may transition into compact phago-lysosome, which then rupture over time, whereas spacious phagosomes remain more stable within the heterogeneous phago-lysosome population. Quantitative image analysis shows a progressive increase in LAMP1 intensity on LAMP1 positive vesicles over time (Supplementary Fig. 1c). This finding is consistent with a previous report highlighting that *Mtb* containing phagolysosome is increasingly marked by LAMP1 with time^35^. Morphometric analysis across time points revealed a persistent heterogeneity among LAMP1-positive vacuoles (Supplementary Fig. 1,2,3.), with both compact and spacious phago-lysosome populations present (Fig1. c, Supplementary Fig. 2b, Movie S1, S2). Direct physical contact between *Mtb* and the phago-lysosomal membrane in both compact and spacious states is suggested, regardless of bacterial viability(Fig. 1f-k) The distinct populations of LAMP1-positive vesicles in both dead and live bacterial infections indicates that just the presence of mycobacterial surface in direct physical contact with the phagosomal membrane is sufficient to induce changes in vesicle architecture ^35^. This data, thus, suggests that changes in lysosomal marker intensity and vesicular structure are not exclusively dependent on bacterial viability or active secretion of effector proteins. Instead, the physical association between *Mtb* and the host phagosomal membrane might serve as a sufficient trigger for these processes, setting the stage for further pathogen- or host-directed modulation and highlighting the role of the bacterial surface^36^.

**Figure 1.**
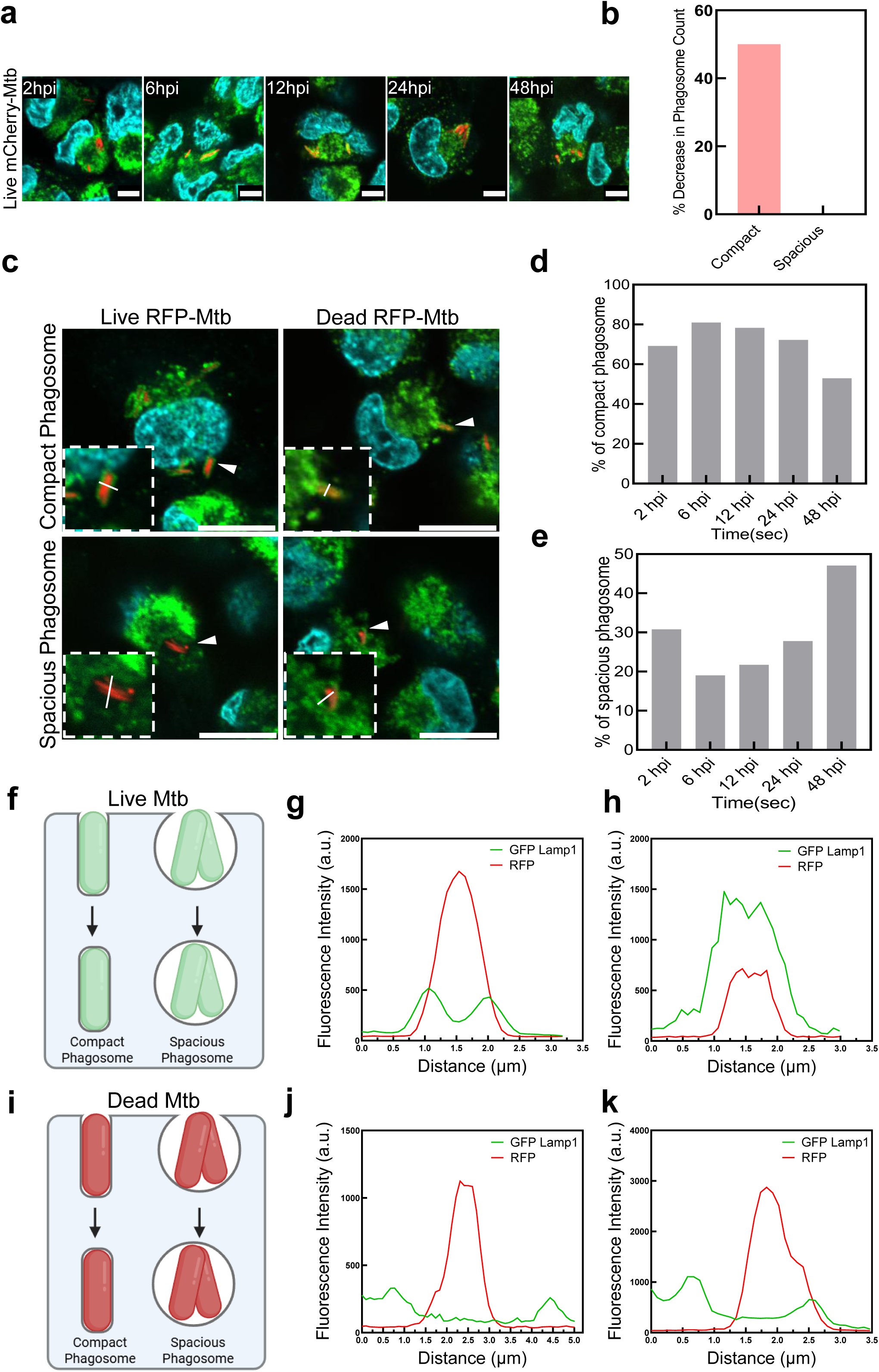
Lamp1 co-localization with *Mtb* increases with time and forms heterogenous populations of compact and spacious phagosomes **a.** Confocal imaging of THP-1 cells infected with Live *Mtb* tagged with *Mtb* tagged with mCherry (red) and immunostained for Lamp1 (green) and fixed with 4% paraformaldehyde/PBS at timepoints-2, 6, 12, 24 and 48hrs post-infection (hpi). Scale bar, 10μm. Representative images of two technical replicates from one independent experiment. (no. of Lamp1 associated *Mtb* = 100). **b.** Comparison of percentage changes in phagosome counts obtained from manual segmentation shows a progressive decline in compact phagosomes up to 48 hpi, while spacious phagosomes remain relatively unchanged. **c.** Representative images for compact and spacious morphology of Lamp1 vesicles encapsulating *Mtb* in infected cells. Scale bar, 10μm. ROI drawn across the structure in ImageJ to retrieve the intensity plot. **d.** Comparison of percentage of compact phagosomes with live *Mtb* obtained from manual segmentation in infected cells till 48hpi. **e.** Comparison percentage of spacious phagosomes and live *Mtb* obtained from manual segmentation in exist in infected cells till 48hpi. **f.** Schematic representation of compact and spacious phagosome in vesicles encapsulating Live *Mtb* in infected cells. **g,h.** Intensity plots of *Mtb* (red) and Lamp1 (green) shows the distribution of fluorescence in compact phagosomes where Lamp1 closely colocalizes with *Mtb* shown by the overlapping of signal peaks. (no. of compact phagosomes = more than 100) **i.** Schematic representation of compact and spacious phagosome in vesicles encapsulating dead *Mtb* in infected cells. **j, k.** Intensity plots of *Mtb* (red) and Lamp1 (green) shows the distribution of fluorescence in spacious phagosomes where Lamp1 and *Mtb* signal peaks show no overlapping. (no. of compact phagosomes = 50)

### Direct contact of mycobacteria induces lipid membrane wrapping and creates local diffusion barriers in the host membrane

To dissect the role of direct contact of bacteria on host phagosomal membrane shaping and remodelling independent of virulence factor like ESAT-6, we used GFP - *M. smegmatis* as a model organism. We used a method to encapsulate live *Mycobacterium smegmatis* inside giant unilamellar vesicles composed of phagosomal membrane lipids (as described in methods and Supplementary Fig. 4a, b). We optimized electroformation to encapsulate live fluorescent bacteria. This method reduces the impact of external impurities such as oil used in emulsion methods and ensures reliable interaction with host membranes. Despite being avirulent, the fast-growing *M. smegmatis* is considered an ideal surrogate for mycobacterial research^37^. We first quantified the viability of the bacteria in the experimental conditions through colony forming unit (CFU) assay and found it to be ∼ 60% (Supplementary Fig. 4c). To capture the dynamics of bacterial interaction in real-time, live bacteria were encapsulated in phagosome membrane mimicking GUVs (40% DOPC, 5.5% DOPE, 22% BSM, 32.5% cholesterol, 0.03% PEG-Biotin) labelled with 0.1% Rhodamine-PE (Rh-PE), immobilized using 0.03% Biotinylated-PEG. Upon monitoring over 60 min, suspended bacteria were found to exhibit random Brownian motion inside the reconstituted phagosome. Interestingly, the area of contact between the bacteria and the host membrane showed an accumulation of host lipids (Fig. 5b, Movie S3). Upon visualizing the GUV in 3D at different planes (Fig. 2c and Supplementary Fig. 5a, b), we observed lipid wrapping over the bacterial surface area as also illustrated in the schematic diagram (Fig. 2a). Such interaction induces membrane bending at the contact area in both live and dead bacterial surface indicating surface interactions at play. Furthermore, fluorescence recovery of Rh-PE at the bacterial contact site is reduced by ∼half following photobleaching in the presence of live bacteria. In contrast, a delayed but full recovery is observed in case of dead bacteria (Fig. 2d), hinting that strong inter-surface interaction between host and live bacteria creates a diffusion barrier. This subsequently leads to lipid segregation within the host membrane. To track the change in lipid reorganization and packaging, we employed two dye model systems (Rh-PE in red and Top-fluor cholesterol in green) and monitored their fluorescence intensity over time (Fig. 2e, Movie S4, Movie S5). We observed that the fluorescence intensity of Top-fluor cholesterol decreases in the vicinity of the contact area (Fig. 2f, h). In contrast, the fluorescence intensity of Rh-PE increases near the contact area, as shown in zoomed images (Fig. 2g, i). Together the live-bacterial direct-contact induces membrane bending, lipid wrapping at the contact area and creates a diffusion barrier resulting in the reorganization of lipids and changes in lipid packing (increased disorderliness) as evident from the marked increase in the fluorescence intensity of Rh-PE in the region ^38^.

**Figure 2.**
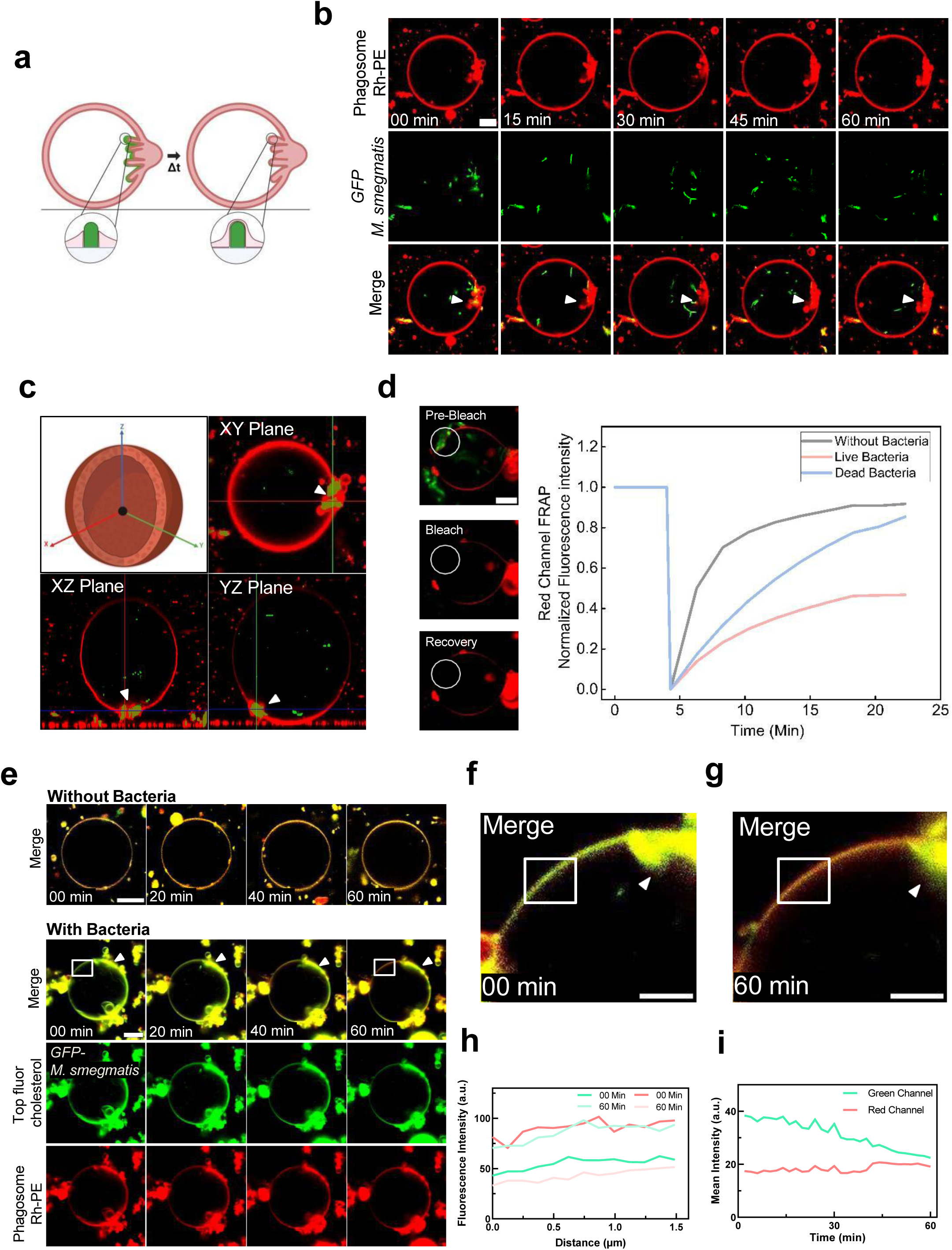
Direct surface contact of *Mycobacterium smegmatis* triggers lipid wrapping and creates diffusion barrier in the phagosome membrane **a.** Schematic representation of the experimental design depicting the impact of direct contact on immobilized GUV mimicking host phagosomal membrane. **b.** Time-lapse confocal imaging of GUVs labelled with 0.1% Rh-PE (red) and immobilized using 0.03% PEG-Biotin encapsulated with live *M. smegmatis-GFP* (green). Scale bar, 10μm. Representative images from three independent experiments. (no. of GUVs = 30-35). **c.** Schematic representation and XYZ Projection of the mycobacteria-mediated curvature change from b. **d.** Mean fluorescence recovery curve after photobleaching experiments of direct contact of bacteria (live and dead) on the membrane. Data points using mean values from three independent experiments. Scale bar, 5μm **e.** Time-lapse confocal imaging of GUVs of phagosomal membrane composition labelled with 0.05% Rh-PE (red) and 0.05% Top Fluor cholesterol (green) and immobilized using 0.03% PEG-Biotin encapsulated with live *M. Smegmatis* (green). Scale bar, 10μm. Images are the representation from three independent experiments. (no. of GUVs = 30-35). Zoomed image of confocal images of region near to direct contact of bacteria at **f.** 0 mins, **g.** 60 mins (Scale bar: 5μm.). White box in e indicates the region for line intensity profile at respective time points. **h.** Fluorescence intensity profile of Topfluor cholesterol (green shades) and Phagosome Rh-PE (Red shades) at two different time points. **i.** Mean fluorescence intensity profile of green channel (Topfluor cholesterol) and red channel (Phagosome Rh-PE).

### Increasing bacterial load induces membrane disorderliness and global phase separation in the host membrane

We next investigated the influence of moderate bacterial load mimicking a spacious phagosome, on the dynamics of the phagosomal membrane. Henceforth, low-to-moderate load and high load mycobacterial conditions are referred to as spacious and compact phagosome mimics, respectively. To capture this, we encapsulated an increased load of bacteria inside the GUV, as illustrated in the schematic (Fig. 3a). We observed that without bacterial load, Rh-PE fluorescence intensity appears to be homogenous throughout the equatorial plane of the membrane. In sharp contrast, in a spacious phagosome mimic, the live bacteria induce striking phase separation in phagosome like compartments as evident from the Rh-PE signal that is restricted to bacterial contact areas (Fig. 3b). However, no significant phase separation was observed in dead bacteria (Supplementary Fig. 6a, b). We observed a change in the fluorescence intensity and thickness of the membrane upon monitoring the GUV mimicking spacious phagosome for 60 minutes (Fig. 3c, Movie S6). Specifically, there is significant mobilization of Rh-PE towards the bottom of the GUV after 60 minutes where the larger pool of bacteria settle down (Fig. 3d, bottom right panel XZ plane). Also, the accumulation of lipids at the bottom of GUVs induces budding at the bottom (Fig. 3d and Supplementary Fig. 7). This suggests that bacteria make the membrane more disordered at the contact areas. Likewise, in both live and dead bacteria condition, the diffusion of Rh-PE on the membrane partially slowed down as evident from the fluorescence recovery after photobleaching at the equatorial plane (Fig. 3e). Super-resolution microscopy revealed that the interactions caused by the bacteria hitting and retracting off the host membrane induced higher disorderliness at the point of contact in the GUVs (Fig. 3f) as seen from the Rh-PE signal at the equatorial plane (Fig. 3g). Interestingly, the membrane sustained the disorderliness despite the bacteria having retracted after hitting the surface and were no more in contact. These observations collectively suggest that increasing bacterial load inside the GUV results in stronger interactions with the host membrane via Brownian motion affecting the lipid packing. This subsequently leads to phase separation and the mobilization of Rh-PE at the bottom as more stable contacts get established as the bacteria settles to the bottom due to gravity. This might also impact the mobility of the lipids at other planes as a result of stretching and lipid reorganization in the membrane.

**Figure 3.**
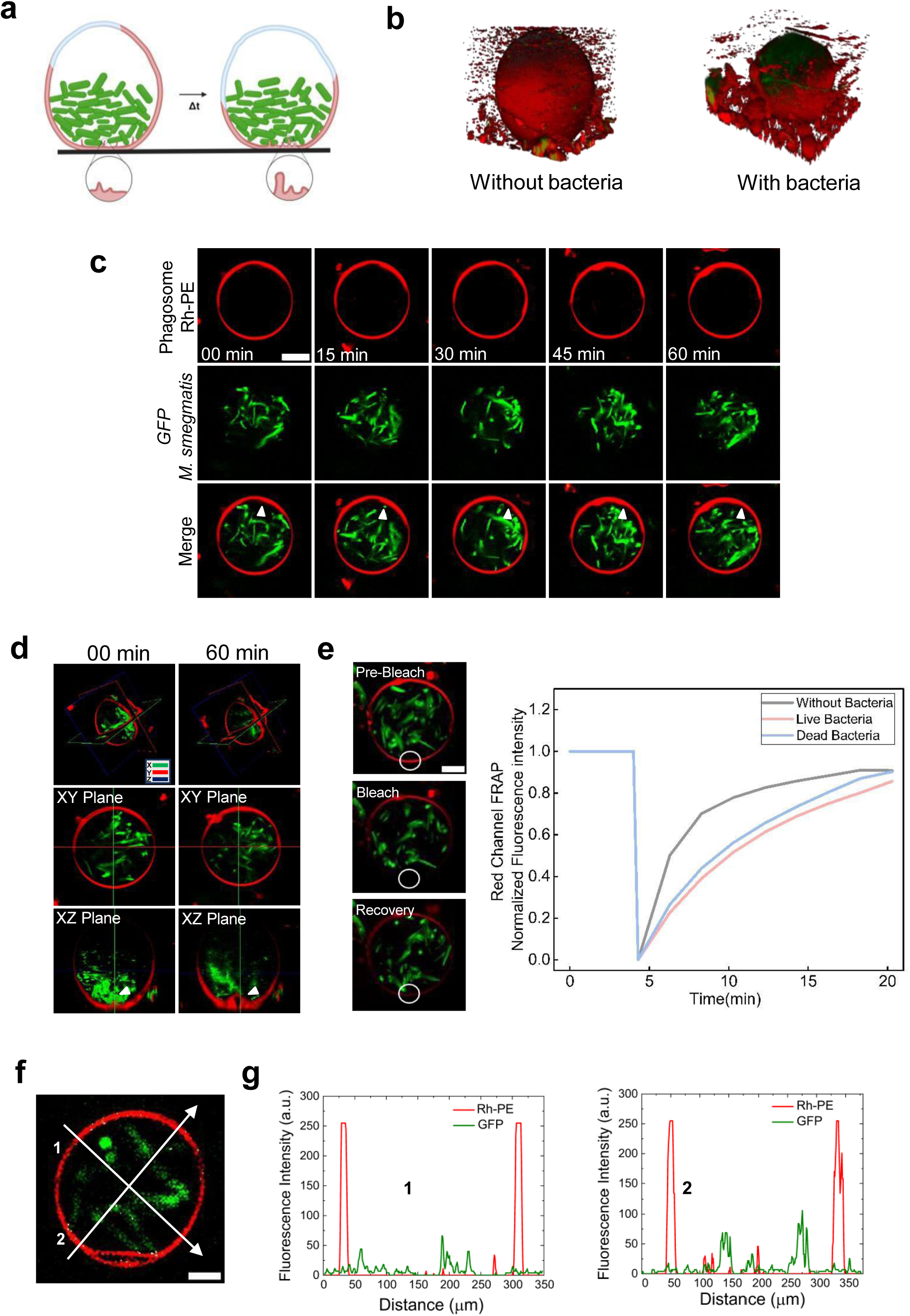
Increasing mycobacterial load induces global phase separation in the phagosome membrane **a.** Schematic representation for the experimental design depicting the impact of a spacious phagosome mimic on immobilized GUV mimicking host phagosomal membrane. **b.** 3D projection of vesicle without and with bacteria exhibiting the bacteria-induced phase separation (no. of GUVs = 30-35). **c.** Time-lapse confocal imaging of GUVs labelled with 0.1% Rh-PE (red) and immobilized using 0.03% PEG-Biotin Biotin mimicking spacious phagosome encapsulated with live *M. smegmatis* (green). Scale bar, 10μm. Representative images from three independent experiments. (no. of GUVs = 30-35). **d.** XYZ Projection of the mycobacteria load mediated changes from c representing the budding event over time. **e.** Mean fluorescence recovery curve after photobleaching experiments of spacious phagosome mimic GUVs on the membrane. Data points using mean values from three independent experiments. Scale bar, 5μm **f.** Structured illumination microscopy (SIM) images of GUVs labelled with 0.1% Rh-PE (red) and immobilized using 0.03% PEG-Biotin encapsulated with moderate load of live *M. smegmatis* (green), 5μm. (no. of GUVs = 30-35). Line intensity plots in **g**. of line 1 and 2 respectively.

### Compact phagosome mimic scaffolds membrane remodelling through bacterial surface-driven condensation

The behaviour of Mycobacteria within the confined space of the mycobacteria-containing vacuole, whether in a compact phagosome or during replication, remains poorly understood, particularly with respect to the establishment of direct contact with the host membrane ^16^. We hypothesized that a substantial increase in mycobacterial load resembling the formation of bacterial clumps within the phagosome ^39^ would induce a higher probability of establishing direct-contact with the phagosomal membrane. To examine the impact of clump formation on the phagosome membrane, we encapsulated a significantly higher load of mycobacteria to mimic the compact phagosome, thereby increasing the contact area with the membrane, as illustrated in the schematic (Fig. 4a). To capture the interactions in real-time, we monitored the compact phagosome mimic for 60 minutes. Striking membrane tampering is induced by the random attachment of the bacteria (Fig. 4b, Movie S7). Interestingly, the host membrane remains permanently bent even in the case where the bacteria tend to retract away from their initial contact area. The bacteria exhibit collective mobility as clumps in a compact phagosome mimic compared to that of the spacious phagosome mimic. In line with previous observations, we noted the accumulation of lipids at the bottom of the membrane after 60 minutes (Supplementary Fig. 9 a, b), and subsequent budding at the bottom of the GUV. The budding phenomenon results from phase separation, where the reorganization and mobilization of liquid-disordered lipids induce localized membrane curvature at contact areas making the membrane more deformable (Supplementary Fig. 9 a, b). To establish phase separation, we used two dye systems as previously described, and observed an intense phase separation of the host membrane by the bacteria (Fig. 4c). Next, to visualize the changes in smaller-sized GUVs similar to phagosome size, we encapsulated 4-6 bacteria within smaller GUVs around size 8-10μm. Remarkably, we found that the bacteria caused complete deformation of the membrane (Supplementary Fig. 9c, d). This observation indicates – for a given bacterial load, a higher contact area ina smaller GUV leads to a significantly greater degree of host membrane deformation compared to larger GUVs. This suggests a size-dependent effect, where the smaller volume and surface area of the GUVs amplify the impact of bacterial interactions, resulting in more pronounced membrane deformations. Super-resolution microscopy reveals the host membrane becomes more disordered at the contact area of the bacteria. The white spots indicate the colocalization point, where the bacteria and host membrane are in contact (Fig. 4d) as evident from the intensity of the overlapping regions of the host and the bacteria (Fig. 4e). Further, the shape of the bacterial clump appears to drive the shape of the host membrane via stretching (Fig. 4f). To investigate the role of bacterial surface on the phase separation of the host lipid membrane, we used a two-dimensional Ising model, where the two states (-1 and +1) represent the lipid ordered and disordered states. Bacterial contacts on the membrane were considered as patches where one kind of lipid has a high probability of accumulating (see Supplementary information for details). We find that when the contacts are few (low bacterial load), global phase separation does not occur. However, the membrane can exhibit local phase separation in the vicinity of the contact site (Fig. 4g). Interestingly, increasing the number of contacts can drive the entire membrane to phase separate (Fig. 4h). The phase separation intensifies as the bacterial contact area on the membrane increases, becoming significantly more pronounced (Fig. 4i). However, increasing the size of bacterial contact on the membrane, keeping the fraction of total area in contact constant, results in a decrease in the degree of phase separation (Fig. 4j-l). This is a consequence of the bacteria-inducing local phase separation. As more bacteria come closer together, there is a significant overlap in the locally phase-separated regions, thereby diminishing the effect on the entire membrane. Bacterial contact-induced phase separation is more pronounced when the lipid interactions are such that the membrane is near the lipid demixing point. (Supplementary Fig. 8a, b, Supplementary text). This suggests that the probability of interaction between bacterial surfaces and the host membrane is influenced by the size of the phagosomal compartment. The disorderliness and degree of phase separation within the membrane are driven by the percentage of bacteria in contact, leading to increased membrane fluidity. As the number of bacteria within a confined volume increase, the probability of interaction and the formation of bacterial clumps also rise. The morphology of the vesicle is determined by the shape of these bacterial clumps (Fig. 3b, f & Supplementary Fig. 9c, d). Further, the fluidized membrane conforms to and wraps around the clumps, resulting in a more rigid scaffold-like structure. Collectively, these concurrent processes are likely to significantly impact membrane parameters.

**Figure 4.**
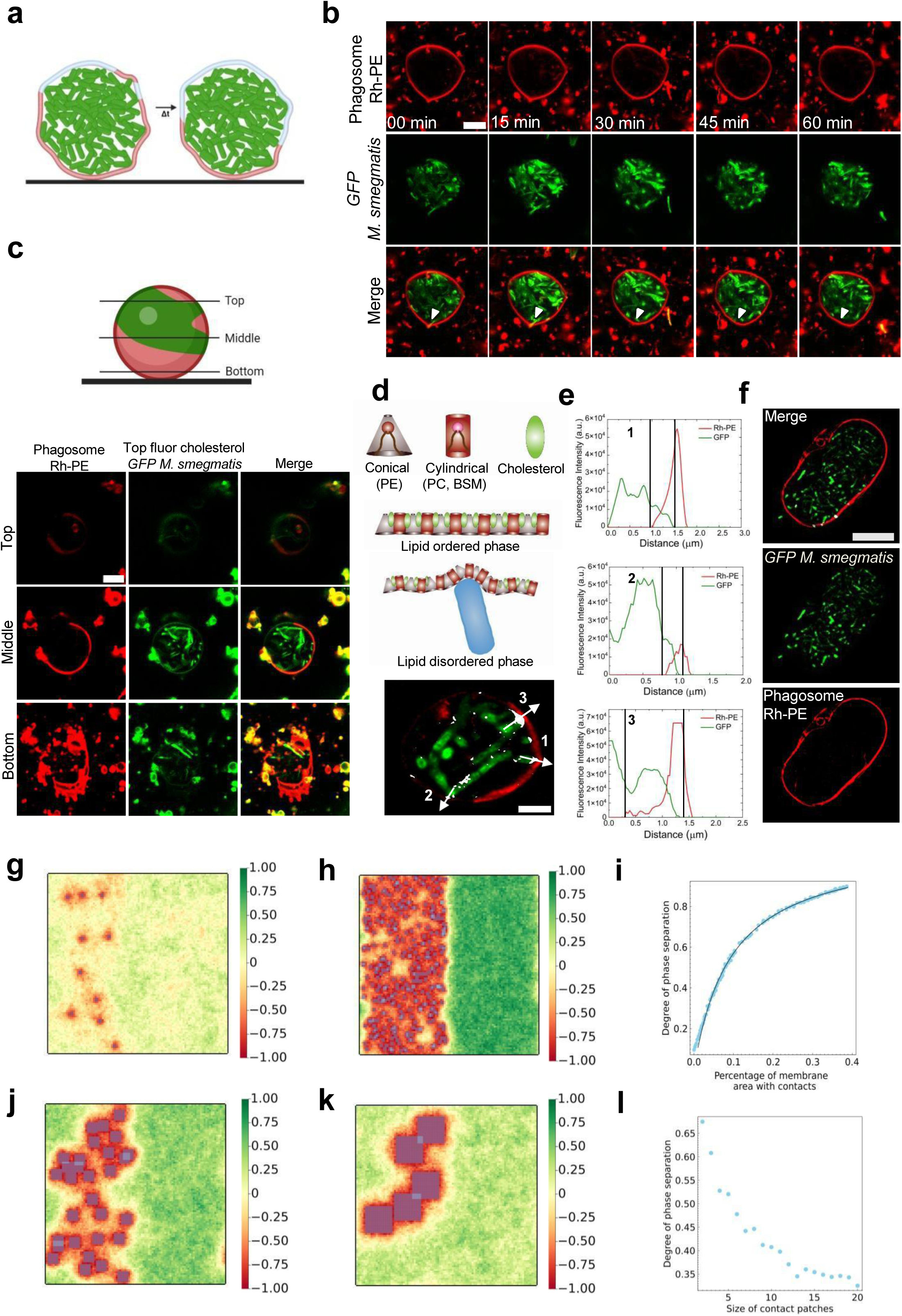
Compact phagosome mimic induces host membrane remodelling through condensation driven by bacterial surface interactions **a.** Schematic representation of the experimental design depicting the impact of a compact phagosome mimic on immobilized GUV mimicking phagosomal membrane. **b.** Time-lapse confocal imaging of GUVs labelled with 0.1% Rh-PE (red) and immobilized using 0.03% PEG-Biotin mimicking Compact phagosome encapsulated with live *M. smegmatis* (green). Scale bar, 10μm. Representative images from three independent experiments (no. of GUVs = 30-35). **c.** Schematic representation and XY plane of the GUV of phagosomal membrane composition labelled with 0.05% Rh-PE (red) and 0.05% Top-Fluor cholesterol (green) and immobilized using 0.03% PEG-Biotin Biotin mimicking Compact phagosome encapsulated with live *M. Smegmatis* (green) representing different planes. Scale bar, 10μm. Images are the representative of three independent experiments. (no. of GUVs = 30-35). **d**. Structured illumination microscopy (SIM) images of GUV, white spot showing degree of colocalization. **e.** Showing line intensity graph plot for Rh-PE (red channel) and GFP (green channel) and marked area with black line denote overlapping region of red and green channel. For the colocalization the points are mentioned in d. **f.** Structured illumination microscopy (SIM) images of GUVs with high bacterial load showing distortion of shape (no. of GUVs = 30-35). Time-averaged state of the membrane. **g.** With a low load, phase separation is observed only near the contacts. Degree of phase separation 0.137 (N_c_ = 10, *p*=2). **h.** With a high load, global phase separation is seen. Degree of phase separation 0.675 (N_c_=300, *p*=2). **i.** Degree of phase separation in the membrane as a function of the percentage of membrane area in contact with the membrane. **j,k,l.** Varying the contact patch size, keeping the fraction of membrane area in contact fixed. j & k show representative time-averaged states for *p*=6 and *p*=15 respectively. (Parameters in all figures: N=100, ϕ=0.5, J*/k_B_T*=0.4, h*/k_B_T*=5.0)

### Mycobacterial load mediated direct contact and ESAT-6 virulence facilitate spacious to compact phagosome switching

We next questioned if there is any correlation between the bacterial load inside the host membrane and the change in the membrane parameters (Fig. 5). To test this, we used micropipette aspiration to aspirate GUVs encapsulating varying loads of the bacteria. The presence of Mycobacteria inside the GUVs was confirmed through bright-field microscopy. Depending on the mobility of the bacteria and shape fluctuation of the GUV, we broadly classified GUVs into Spacious phagosome mimic and Compact phagosome mimic (Fig. 5a-c, Movie S8, Movie S9, Movie S10). Spacious phagosome mimic seems to exhibit higher membrane fluctuations and bacterial motion compared to Compact phagosome mimics and phagosome mimic (i.e., without bacterial load) (Fig. 5a-c, Movie S8, Movie S9, Movie S10). Simultaneously, the geometry (aspiration length *L^p^* and vesical radius *R_v_*, see Fig. 5a, b, c) of the GUV was monitored to document changes in membrane parameters by the interaction of bacteria with the host membrane. Increasing the bacterial load inside the GUV results in a higher probability of establishing bacterial contact with the host membrane as seen in Fig. 3, 4. To quantify these changes in membrane properties, we aspirated Spacious phagosome mimic with a constant change in pressure (i.e., *ΔP= 50 Pa*). We observed the aspiration length *Lp* increases significantly in bacteria loaded spacious phagosome as compared to phagosome mimic without bacteria (Fig. 5a, b, d). Spacious phagosome mimic is found to reduce the area stretch modulus by 5 folds as compared to the phagosome mimic (Fig. 5e). Likewise, the bending modulus of the host membrane is reduced by 6 folds as compared to the phagosome mimic (Fig. 5f). Surprisingly, no change in L_p_ is observed for the Compact phagosome (Fig. 5c, d). The aspiration length (*L_p_* = 0) suggests an absence of direct area expansion, precluding the calculation of both area stretch modulus and bending modulus. Interestingly, significant deposition of host membrane lipid is observed over the surface of the pipette (Fig. 5c, Movie S10). This observation suggests that as the bacterial load increases, the fluidity of the host membrane also increases, leading to greater membrane deformability. In contrast, under high load conditions, bacteria aggregate to form a scaffold-like surface. The fluidized host membrane, influenced by bacterial activity, then wraps around this clump. This interaction collectively results in increased membrane rigidity. These bacterial interactions are efficient in remodelling the membrane properties but are not sufficient for the bacteria to escape from the phagosome membrane.

Furthermore, to investigate the role of virulent secretory proteins in bacterial escape, we treated these three conditions with a low concentration (2.5 μM) of monomeric ESAT-6 protein and monitored them for 2 hours. Interestingly, we observed that in this low concentration and the absence of bacteria, there was no significant change in the membrane shape (Fig. 5g, Movie S11). However, spacious phagosome significant budding in the membrane was observed after 60 minutes (0.0667 FPS) (Fig. 5h, Movie S12). As observed in the previous experiment, under a Compact phagosome, the GUV membrane became more rigid compared to the spacious phagosome (Fig. 5c), creating favourable conditions for membrane rupture and bacterial escape from the phagosome. Subsequent treatment with ESAT-6 induces phagosome membrane rupture resulting in bacterial escape (Fig. 5i, Supplementary Fig. 10 a, b, Movie S13). This indicates that bacterial load enhances the membranolytic activity of the virulence factors. The proposed model suggests that direct contact with Mycobacteria affects the maturation of the Mycobacteria-containing vacuole (MCV). This interaction reduces the bending rigidity of the MCV membrane, that might alter the kinetics of maturation machinery, besides enhancing ESAT-6-mediated membrane disruption in a bacterial load-dependent manner.

**Figure 5.**
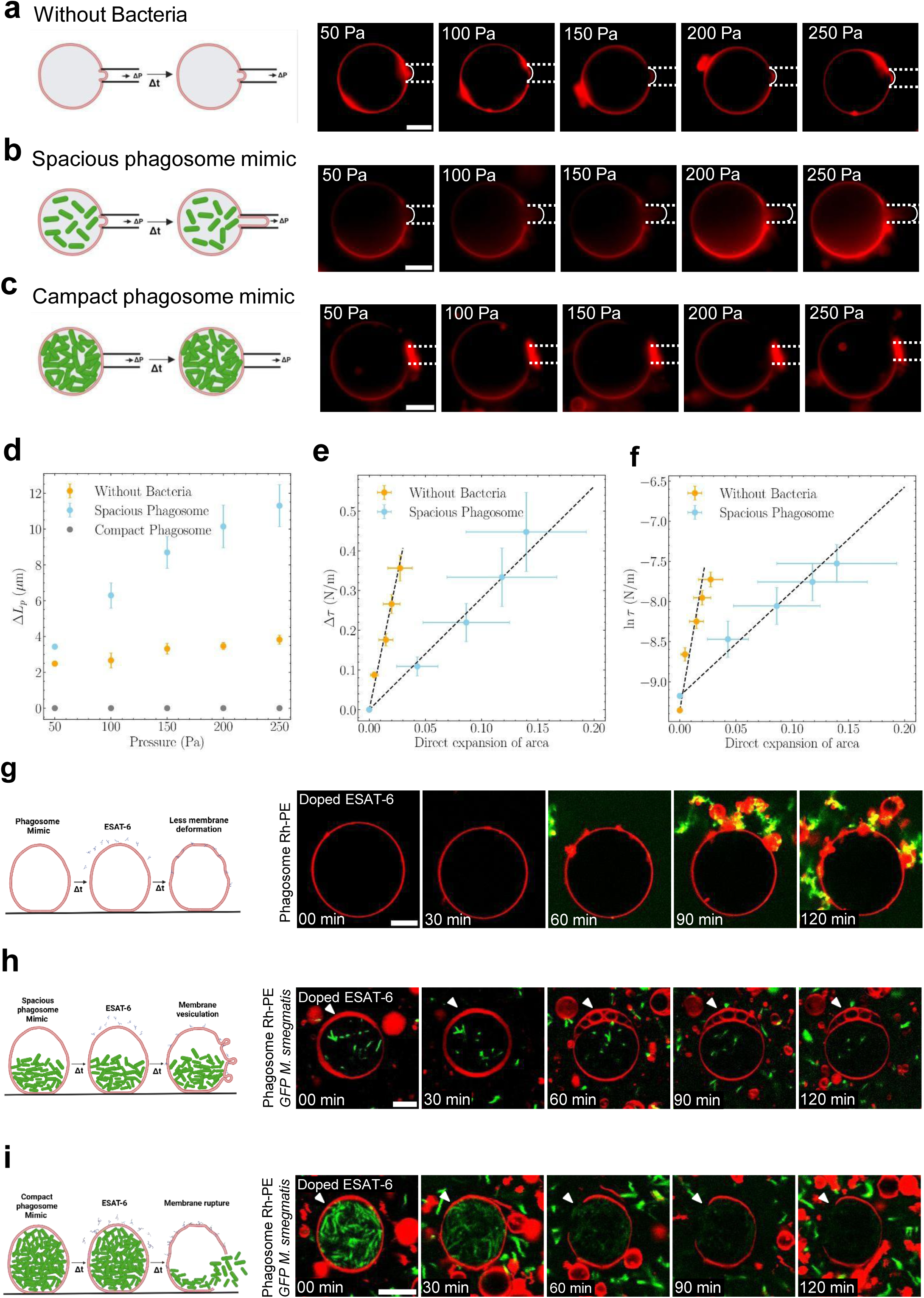
Direct surface contact of mycobacteria enhances the virulence of ESAT-6 and facilitates spacious to compact phagosome switching **a.** Schematic representation and time-lapse Epi-fluorescence imaging of GUVs labelled with 0.1% Rh-PE (red). Scale bar, 10μm. Aspirated with Pressure (Δp = 50Pa) and monitored over time. Representative images from three independent experiments. **b.** Schematic representation and time-lapse Epi-fluorescence imaging of GUVs labelled with 0.1% Rh-PE (red) mimicking spacious phagosome with live *M. smegmatis* (green). Scale bar, 10μm. Aspirated with Pressure (Δp = 50Pa) and monitored over time. Representative images from three independent experiments. **c.** Schematic representation and Time-lapse Epi-fluorescence imaging of GUVs labelled with 0.1% Rh-PE (red) mimicking compact phagosome with live *M. smegmatis* (green). (green). Scale bar, 10μm. Aspirated with Pressure (Δp = 50Pa) and monitored over time. Representative images from biologically three independent experiments. **d.** Plot for the length of protrusion(ΔL_P_) vs pressure (P) from a,b and c. Data points shown are the Means ± SD of three independent experiment. **e.** Quantifying the area stretch modulus(k_a_) from the change in tension(Δτ) vs direct expansion of area plot from a, b and c. Orange and blue spots with bars: experimental points for without bacteria and Spacious phagosome, respectively, average ± SD. Dotted line: Linear fit, y= a*x+b, for without bacteria a = 13.5 ± 1.09, b = (-1.25 ± -0.49)*10^-6^, R^2^ = 0.125 and for Spacious phagosome a = 2.81 ± 0.10 b.(-1.81 ± 0.727)*10^-7^, R^2^ = 0.056455, K_a(without bacteria)_ = 13.5 ± 1.09mN/m K_a(Spacious phagosome)_ = 2.81 ± 0.10 mN/m **f.** Quantification of the bending modulus (k_b_) from the semi-log plot of tension versus direct expansion of area from a, b, c. Orange and blue spots with bars: experimental points for without bacteria and Spacious phagosome, respectively, average ± SD. Dotted line: Linear fit y= a*x+b, for without bacteria a = 81.52 ± 1.489, b = -2.45 ± (6.74*10^-6^), R^2^ = 0.605 and for Spacious phagosome a = 13.021 ± 0.9696, b = -2.27 ± (4.28*10^-7^), R^2^ =-0.097, K_b(without bacteria)_ =(1.33 ± 0.022)*10^-20^ J, K_b(Spacious phagosome)_ = (0.21±0.02)*10^-20^ J **g.** Time-lapse confocal imaging of GUVs labelled with 0.1% Rh-PE (red) and immobilized using 0.03% PEG-Biotin incubated with 2.5μM doped FITC-ESAT-6 (green). Scale bar, 10μm.Images are representative of 3 independent experiments (no. of GUVs = 30-35). **h.** Time-lapse confocal imaging of GUVs labelled with 0.1% Rh-PE (red) and immobilized using 0.03% PEG-Biotin encapsulated with moderate load of live *M. smegmatis* (green) and incubated with 2.5μM doped FITC-ESAT-6 (green). Scale bar, 10μm.Images are representative of 3 independent experiments (no. of GUVs = 30-35). **i.** Time-lapse confocal imaging of GUVs labelled with 0.1% Rh-PE (red) and immobilized using 0.03% PEG-Biotin encapsulated with high load of live *M. smegmatis* (green) and incubated with 2.5μM doped FITC-ESAT-6 (green). Scale bar, 10μm (no. of GUVs = 10-15).

**Figure 6.**
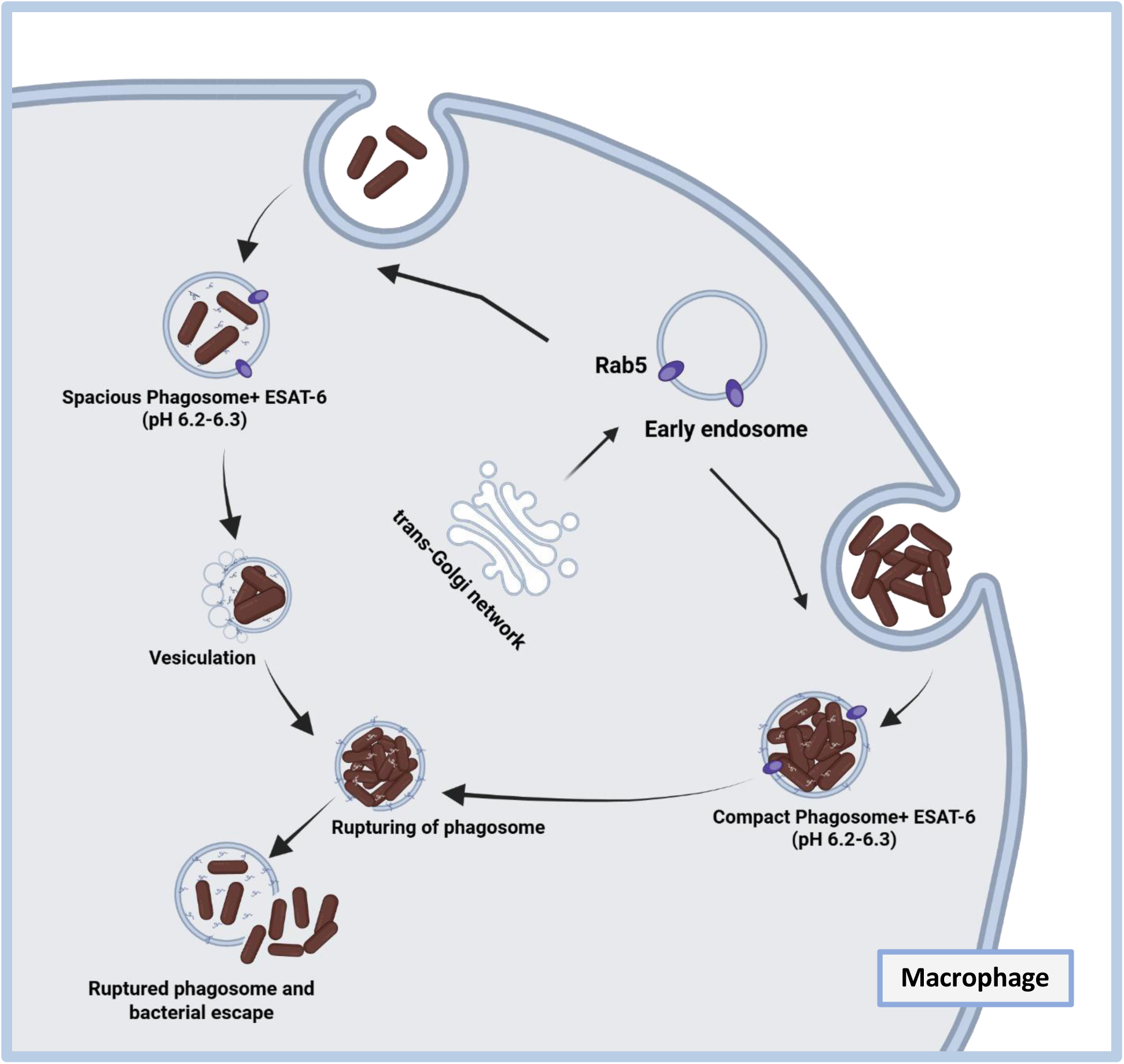
Schematic representation of the proposed mechanism of direct contact of mycobacteria enhances the virulence of ESAT-6

## Discussion

Many intracellular pathogens, including *Mycobacterium tuberculosis* (*Mtb*), manipulate host membranes via secreted effector proteins. However, the role of direct physical contact between bacteria and host membranes and the mechanisms that dictate the pathogen residency time remain unknown^1,40^. Here, we show that bacterial internalization is highly dynamic and heterogeneous, with both compact and spacious LAMP1-positive vacuoles persisting over time, independent of bacterial viability (Fig. 1). Previous studies indicate that internalization is governed not only by host or pathogen specificity but also by the degree of interaction ^41^. For *Mtb*, the extent of interaction influences intracellular uptake and contributes to heterogeneous compartmentalization^42–44^. Our findings reveal that *Mtb* exploits direct physical contact with host phagosome membranes to mediate switching from spacious to compact phagosomal state (Fig. 1b,d,e). We demonstrate that this transition is governed by membrane remodelling by bacteria, where bacterial load and contact area serve as determinants of phagosome fate. At low-to-moderate bacterial loads mimicking spacious phagosome-like compartments, a significant reduction in both bending and area stretch moduli is observed accompanied by membrane fluidization and vesiculation. This process initiates when mycobacterial contact induces membrane bending, lipid wrapping and reorganization at the membrane interface, creating diffusion barriers that promote phase ^46^ (Fig. 2). Notably, these effects are amplified when host membranes are near their lipid lipid demixing point. The change in the Rh-PE and Top-Fluor Cholesterol signal suggests that the bacteria may be extracting cholesterol from the host membrane as *Mycobacteria* are known to scavenge cholesterol from the host for their metabolism ^45^. Indeed intracellular vacuolar compartments represent soft confinements that can be subjected to remodelling and deformation by localized forces such as the active propulsion by motile bacteria^46^.

The extent of host membrane phase separation depends on bacterial load (Fig. 2-4). Under spacious phagosome mimics, freely moving bacteria contacting the membrane at multiple planes induced fluidization and disordered protrusions (Fig. 3d,e). In contrast, compact phagosome mimics with higher bacterial load cause global reshaping and complete phase separation driven by bacterial clumping (Fig. 4b). The phagosome membrane’s proximity to a lipid demixing point further enhances phase separation upon bacterial contact (Fig. 3b). Indeed, previous studies ^47–51^ have suggested that cell membranes lie close to the lipid demixing point^52^. Bacterial surface may induce condensation of host membrane interfaces that are close to the lipid demixing point driving phase separation. Such a phenomena might also explain the lipid zippering proposed for bacterial surface lectin LecA with the host cell glycosphingolipid Gb3 sufficient for driving complete internalization of the *Pseudomonas aeruginosa* ^5^. Self-propelling motile bacteria are known to induce stronger non-equilibrium fluctuations in the membrane, resulting in the formation of long protruding tubes ^30^. Unlike motile bacteria, host membrane deformation by mycobacteria is driven by inter-surface interactions and dictated by the ratio of available surface area of the host membrane and the area of direct-contact. Importantly, this mechanical priming synergizes with ESAT-6 activity, as the softened membrane becomes more susceptible to the effects of the virulence factor. Load dependent shape deformations of encapsulated giant vesicles have also been observed for self-propelled janus colloids that induce highly branched tether-like protrusions at low/moderate loads and globular reshaping at higher loads capturing vesicle shapes that do not exist in equilibrium systems ^53^.

The striking change in the area stretch modulus and the bending modulus of the compartmental membrane with respect to the bacterial load not only renders it more deformable but also might influence the dynamics of phagosome maturation. Under spacious phagosome mimic, ESAT-6 activity induces membrane vesiculation, reducing the available area of the host membrane (Fig. 4h), that may eventually progress into a compact phagosome mimic (Fig. 5i). This ultimately leads to phagosome rupture, similar to the effects observed under high bacterial load conditions (Fig. 5i). Vesiculation may represent a transient intermediate in the conversion from spacious to compact phagosomes. Such compact phagosome compartments preferentially facilitate the activity of secreted virulence factors, thereby promoting *Mtb* access to the host cytosol ^44^. ESAT-6 has been reported to induce independent events like vesiculation and leakage ^54^. The ESX-1 secretion system facilitates direct interaction with host membranes^23^, and targeting sterol-rich microdomains to disrupt the Mycobacterium-Containing Vacuole (MCV)^55^. The bending modulus and tension of the membrane are known to regulate the binding kinetics of proteins, the shape generation of the membrane during endocytosis as well as the rate of budding for inter-organelle trafficking ^56–58^. The direct-contact mediated manipulation of the phagosomal compartment could facilitate ESAT-6-mediated disruption while potentially delaying phagosome maturation. Indeed, the binding of Ras family proteins is known to be sensitive to membrane tension ^56, 59^. The mechanical properties of the membranes are critical for processes involving fission and fusion of membranes ^57, 60, 61^. Together, we propose that the ability to switch from spacious to compact states may represent an evolved strategy allowing *Mtb* to either persist within protective compartments or escape into the cytosol when conditions favor rupture regulating their intracellular residency time.

## Material & Methods

### Materials

1,2-Dioleoyl-sn-glycero-3-phosphocholine (DOPC), 1,2-dioleoyl-sn-glycero-3-phosphoethanolamine (DOPE), BSM, di-stearoyl phosphatidyl ethanolamine-PEG (2000)-biotin, and cholesterol were obtained from Avanti Polar Lipids. The fluorescent lipid lissamine rhodamine B sulfonyl (Liss Rhod PE) and Top-fluor cholesterol were also sourced from Avanti Polar Lipids. Avidin-coated chamber slides were acquired from Sigma-Aldrich.

### Preparation of GUV

Giant unilamellar vesicles (GUVs) that imitate the phagosomal membrane were made up of 40% DOPC, 5.5% DOPE, 22% BSM, 32.5% cholesterol, 0.03% PEG-Biotin, and 0.1% Rhod PE. GUVs were created by getting and slightly altering the protocol given in^57^. In summary, conductive ITO-coated glass from Nanion Technologies, GmbH was equally covered with 15μL of the lipid mixture at a concentration of 5 mg/ml, and it was vacuum-dried for a minimum of three hours. After rehydrating the dried lipid film in 300 ± 5 mOsm sucrose solution of pH (6-6.5), the GUVs developed for three hours at 40-45°C with a sine voltage of 2V and 10 Hz. Using 7H9 media containing precultured Mycobacteria, the lipid film was rehydrated in order to encapsulate live Mycobacteria within the GUVs.

For encapsulation experiments, *Mycobacterium* cultures were grown in 7H9 medium to an optical density OD of 1.0–1.2. The bacterial suspension was subsequently used for encapsulation within giant unilamellar vesicles (GUVs). To assess bacterial viability under the experimental conditions, colony-forming unit (CFU) assays were performed, which indicated that approximately 60% of the encapsulated bacteria remained viable.

### Microscopy

An 8-well Ibidi chamber slide was coated with 10μl of 1 mg/ml streptavidin, followed by the addition of 200μL of biotinylated GUVs. The GUVs were then incubated for 15 minutes to cause immobilisation. Leica TCS SP8 LCSM imaging was carried out with the proper Rhodamine-PE (561 nm) and Alexa-488 (488 nm) laser. Every experiment was conducted using the same laser power and gain settings.

Initially, images were taken with a lower laser intensity for the Fluorescence Recovery after Photobleaching (FRAP) measurements. A 5μm radius of bleaching was achieved within the chosen ROI after 30 seconds of photobleaching at maximum laser power. The recovery curve was then measured, and the laser intensity was adjusted to the attenuated level. The GUV under visualisation was photobleached at its equatorial plane. Following three repetitions, the FRAP curves were standardised for each condition.

### Expression and Purification of ESAT-6

The ESAT-6 protein was expressed using the Escherichia coli BL21 (DE3, pLysS) strain. The strain was transformed with the plasmid pET22b containing the *Mycobacterium tuberculosis* gene Rv3875, which encodes the ESAT-6 protein. Transformed cells were grown in Luria-Bertani (LB) medium at 37°C until the optical density at 600 nm (OD600) reached approximately 0.6. Protein expression was induced by adding 1 mM isopropyl β-D-1-thiogalactopyranoside (IPTG), followed by overnight incubation at 18°C. Post-induction, the bacterial cells were harvested by centrifugation and lysed via sonication. The cell lysate was then subjected to affinity purification using nickel-nitrilotriacetic acid (Ni-NTA) resin, per the manufacturer’s instructions (GE Healthcare, UK). The eluted fractions containing the His-6-tagged ESAT-6 protein were analysed using SDS-PAGE to confirm protein purity. Purified fractions were subsequently dialyzed against an appropriate buffer, concentrated, and stored at -80°C in aliquots for further use.

### Micropipette Aspiration

Using a 100X objective lens, all measurements were made using an Olympus IX83 epifluorescence microscope fitted with Hoffman modulation optics. Three-axis hydraulic micromanipulators are utilised to position the pipette. Capillaries with a diameter of 10μm are used to extract micro-pipettes (Narashige Micro Forge MF-900, Sutter Instruments CO P-87). On top of the unfinished DIY coverslip, 100 μl of GUVs were put. The suction pressure P applied to a bilayer vesicle produces a uniform membrane tension (τ), which is described by a geometric relation based on the pipet (Radius)R_pipette_ and vesicle (Radius)R_GUV_ of the vesicle spherical segment exterior to the pipette,

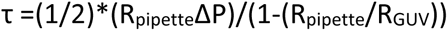

^62, 63^. Movement of the projection length L_p_ inside the pipette directly measures the area increase. Using measurements of the vesicle spherical segment and cylindrical projection dimensions, we numerically computed changes in apparent area Δa for each displacement ΔL_p_ of the projection length with simple geometric formulae and the constraint of a fixed vesicle volume ^64^. The proportionality between apparent area and length is easily seen in the following first-order approximation:

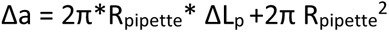

While the reference area a_0_= 4π*R_GUV_^2^ corresponds to a spherical vesical radius R_GUV._ These formulas combine to give the following expression for the areal strain:

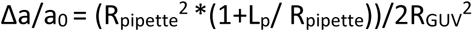

Displacement of the projection length of an aspirated vesicle under a change in suction pressure is shown in Fig.4d. Even though the apparent area can be measured over a tension range from 76 μN/m to 550 μN/m as demonstrated in Fig. 4e to determine the area stretch modulus (K_a_), we plot the change in tension (τ) against direct area of expansion, and the resulting slope is K_a._to determine the bending modulus (K_b_), we plot the semi-log of tension(τ) against the direct area of expansion, and resulting slope is K_b._ All the experiments were performed at room temperature^64^.

### Image Analysis

Image analysis and processing were done using the ImageJ program. The average fluorescence intensity was quantified using the line intensity profiling at the direct contact sites along the equatorial plane of GUVs. The same detection configuration was followed for every experiment. Every experiment set used the same detection configuration. ImageJ was used to manually quantify the diameters of the vesicles.

### Mycobacterial strains and growth conditions

*M*. *tuberculosis* H37Rv expressing mCherry strains were grown in Middlebrook 7H9 (BD Difco 271310) supplemented with 10% OADC (BD Difco 211886) at 37 °C. Before infection, bacterial clumps were removed by centrifugation at 80 × g and supernatant was pelleted, re-suspended in RPMI media, and used for infection, as discussed previously ^65^. All the experiments were performed inside a BSL-3 facility.

### Cell culture and infection

THP-1 monocytes were cultured in RPMI 1640 (Gibco™ 31800022) supplemented with 10% fetal bovine serum (Gibco™ 16000-044). THP1 monocytes were differentiated to macrophages by treatment with 20 ng/ml of phorbol myristate acetate (PMA) (Sigma-Aldrich, P8139) for 24 h followed by incubation in PMA-free RPMI 1640 medium for 24 hours and used for infections. THP-1 monocyte-derived macrophages were incubated with mycobacteria at an MOI of 10 for 2 h followed by the removal of extracellular bacteria by multiples washes. Infected cells were fixed with 4% paraformaldehyde (Sigma 158127) at 2, 6, 12, 24 and 48 hpi and were used for subsequent experiments.

### Immunostaining, imaging, and image analysis

For immunostaining of LAMP1 marker, differentiated THP-1 macrophages after infection were fixed with 4% paraformaldehyde, washed with PBS, and permeabilized with SAP buffer (0.2% saponin (Sigma-Aldrich S4521), 0.2% gelatin (Himedia Laboratories TC041) in PBS) for 10 min at room temperature. Primary antibody Lamp1 (DSHB, clone H4A3) was prepared in SG-PBS (0.02% saponin, 0.2% gelatin in PBS) and incubated at room temperature for 2 hours. After washing with SG-PBS, cells were incubated in AlexaFluor 647 (Life Technologies, Invitrogen) prepared in SG-PBS for 1 h at room temperature, washed, stained with 1 μg/ml of 4[prime],6-diamidino-2-phenylindole (DAPI) for 30 minutes at room temperature. The coverslips were then mounted using a Mowiol-based mountant and imaged using Olympus FV3000 confocal microscope using a 60x objective.

Images were analyzed by CellProfiler 4.2.8. and pipelines similar to those previously established^66^ were used. In all cases, images were segmented to identify nuclei, cells, bacteria, and Lamp1-positive compartments. Objects such as bacteria and Lamp1-positive vesicles were related to individual cells to obtain single cell statistics, and multiple parameters relating to their numbers, sizes, intensities, as well as *Mtb*-Lamp1 associations were extracted. MS Excel and GraphPad 8.0.1 were used for data analysis and plotting graphs. Statistical significance between different sets was determined using Mann-Whitney unpaired test *t* test with unequal variance.

## Acknowledgements

We thank Rohan Dhiman for the kind gift of the ESAT-6 plasmid and GFP-*M. smegmatis* strain. We thank Shubhay Arvind Dikkar and Sajag Kumar for their inputs during technical discussion. We also thank Abik Hameem P.M. and Dipam Naskar for their initial help with micropipette experiments. We acknowledge all the members of the M.S. lab for critical reading of the manuscript. We thank the Biosafety and Central Imaging and Flow Cytometry (CIFF) facilities at NCBS. SC is supported by the Ramalingaswami Re-entry Fellowship (BT/HRD/35/02/2006), Ministry of Science and technology, Government of India. MS gratefully acknowledges the DBT-Wellcome Trust India Alliance (Grant-IA/I/20/2/505212) and Department of Atomic Energy for financial support. VS acknowledges core funding from NCBS-TIFR and Department of Atomic Energy.

