## Supplementary data for "Mycobacterial load and direct contact primes spacious to compact phagosome switching for egress"

### Supplementary Figures

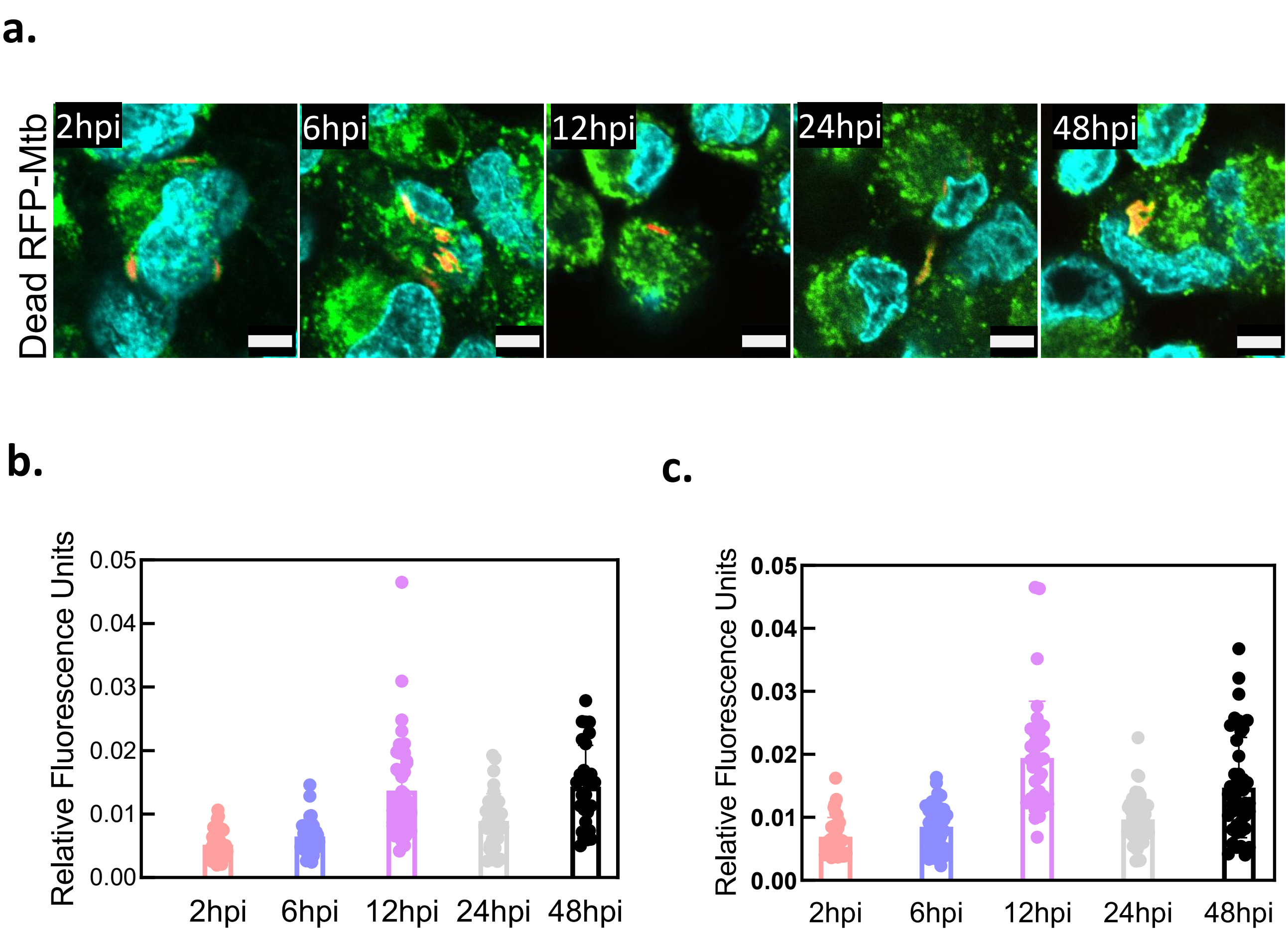


Supplementary Figure 1.

**a.** Confocal imaging of THP-1 cells infected with dead (Heat Killed) *Mtb* tagged with mCherry (red) and immunostained for Lamp1 (green) and fixed with 4% paraformaldehyde/PBS at timepoints- 2, 6, 12, 24 and 48hrs post-infection. Scale bar, 10µm. Representative images of two technical replicates from one independent experiment. (no. of Lamp1 associated *Mtb* = more than 100). **b.** Quantitative analysis of Lamp1 associated dead (Heat Killed) *Mtb* across time showing increasing Lamp1 recruitment and colocalization with bacteria. Statistical significance calculated by Mann-Whitney unpaired test *t* test with unequal variance. **c.** Quantitative analysis of Lamp1 associated Live *Mtb* across time showing increasing Lamp1 recruitment and colocalization with bacteria. Statistical significance calculated by Mann-Whitney unpaired test *t* test with unequal variance**.**

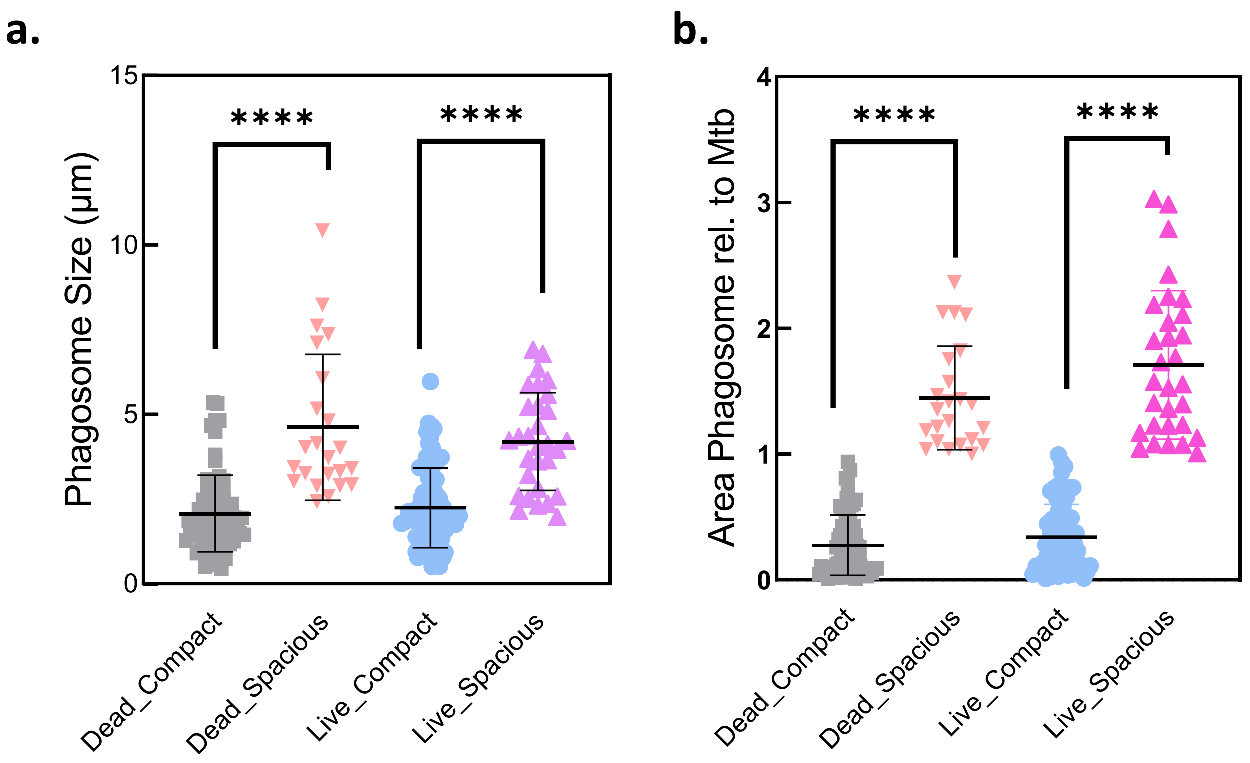


Supplementary Figure 2.

**a.** Comparison of the areas of phagosomes and *Mtb* (live and dead) obtained from manual segmentation shows that two distinct populations of compact and spacious phagosomes exist in infected cells till 48hpi. **b.** Comparison of the relative areas of phagosomes and *Mtb* (live and dead) obtained from manual segmentation shows that two distinct populations of compact and spacious phagosomes exist in infected cells till 48hpi.

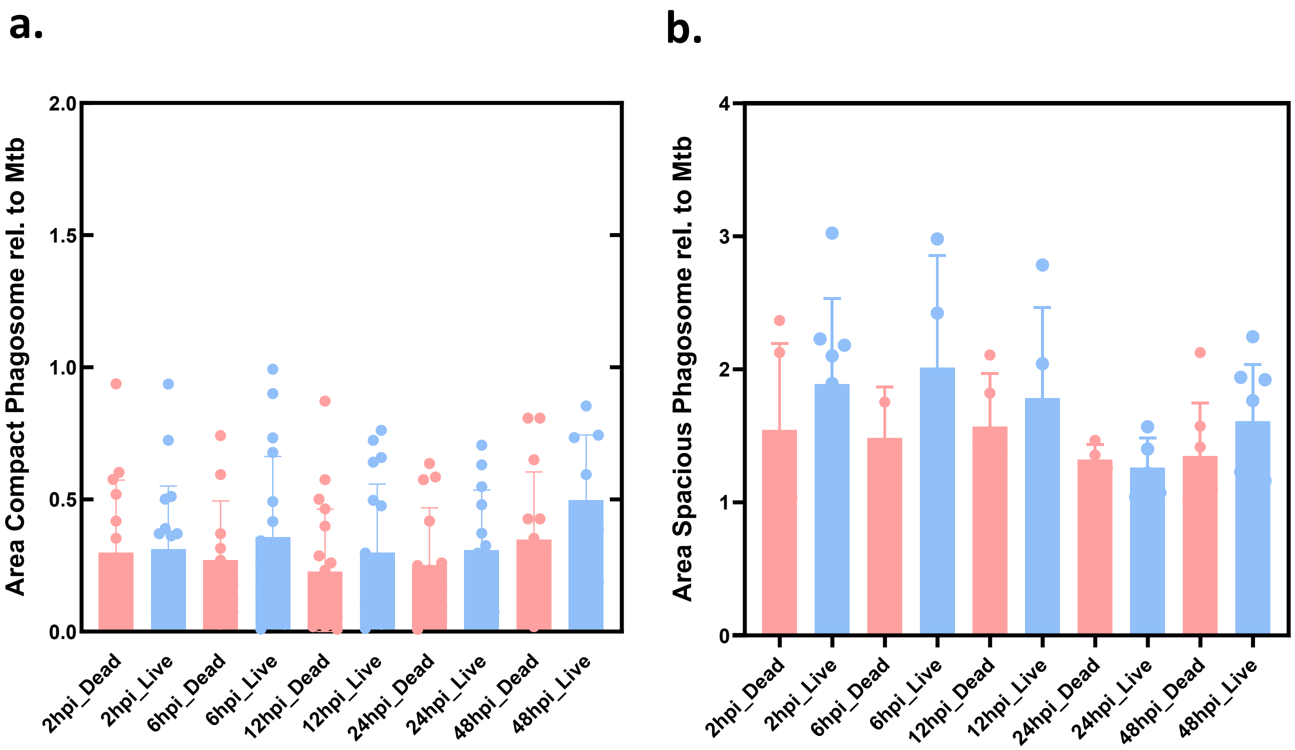


Supplementary Figure 3.

**a,b:** areas of compact and spacious phagosomes relative to *Mtb*, both live and dead, obtained from manual segmentation across time points 2, 6, 12, 24 and 48hpi.

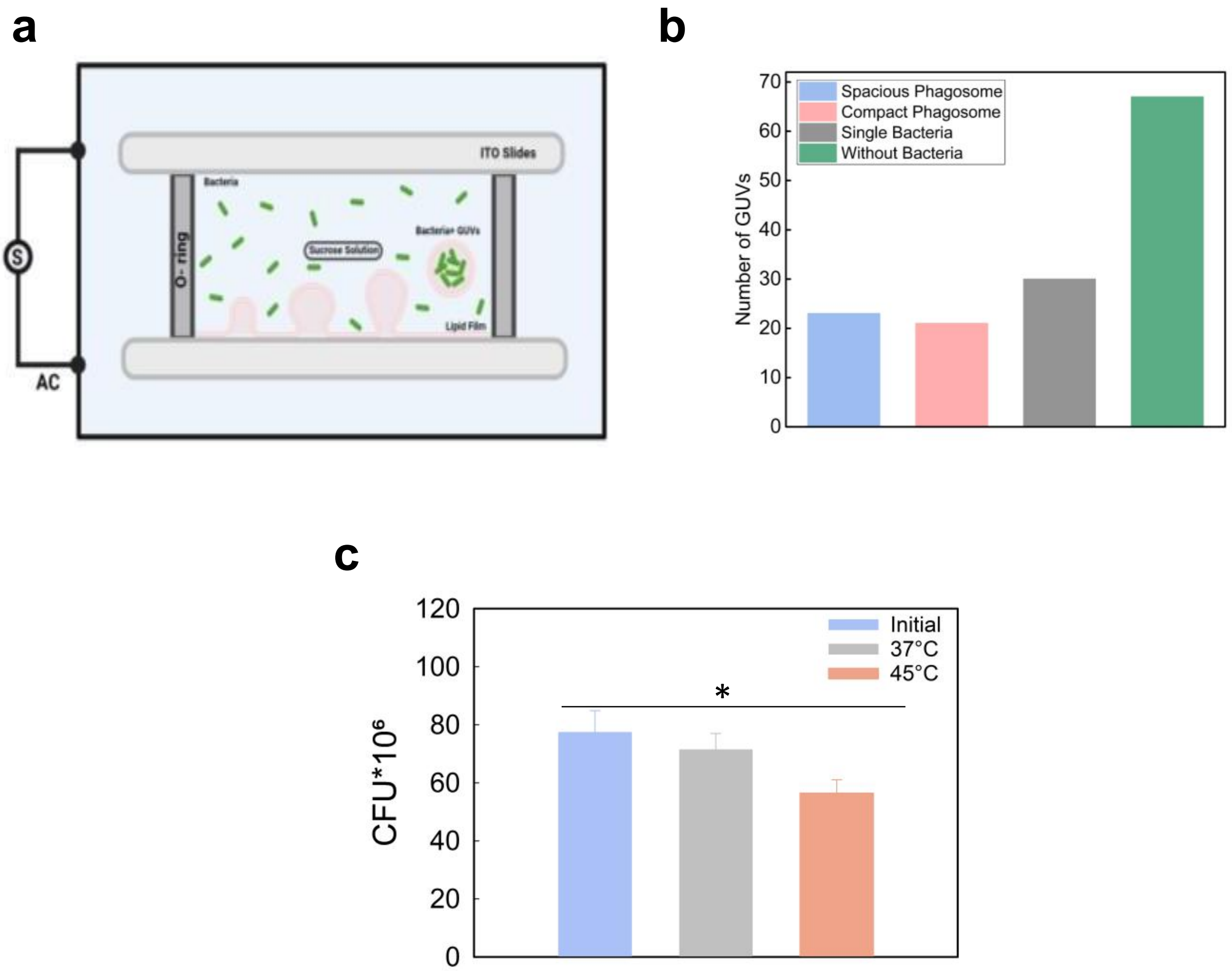


Supplementary Figure 4.

**a.** Schematic representation of experimental design depicting electroformation and encapsulation of bacteria inside the GUV mimicking the phagosomal membrane. **b.** Statistical analysis of the encapsulated bacteria at variable load conditions (no. of GUVs = more than 100). **c.** Viability of the encapsulated *Mycobacterium smegmatis* quantified by colony-forming units at different temperatures to mimic conditions similar to electroformation. Data shown as Mean ± SD. (*$p<0.005$ in one-way ANOVA).

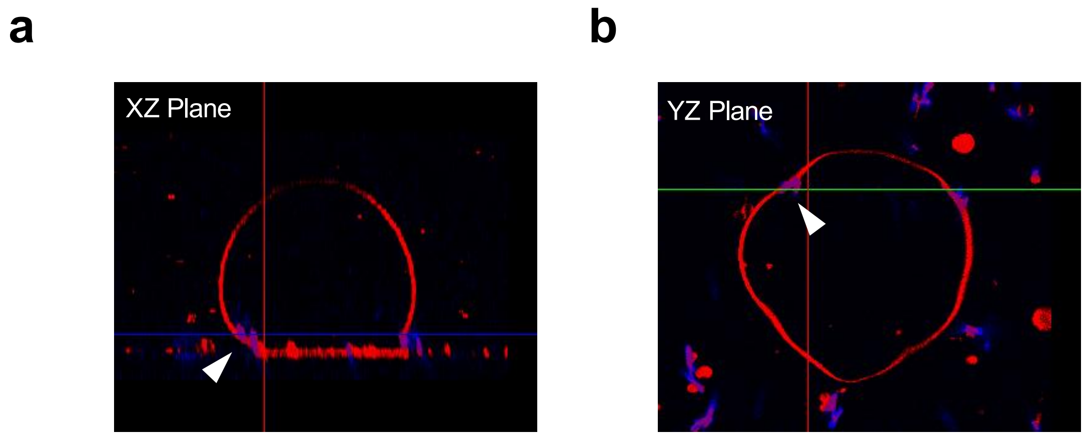


Supplementary Figure 5.

**a and b** shows the XYZ projection of the dead *M. smegmatis*(stained with DAPI) mediated curvature change.

a b

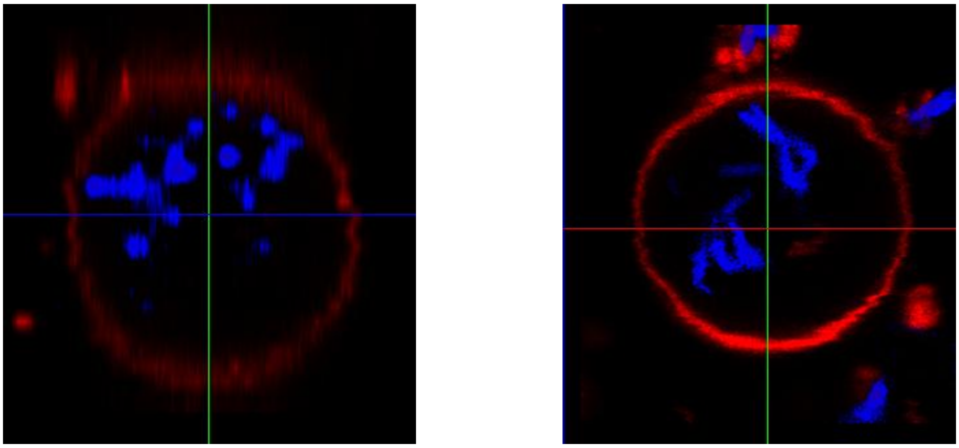

Supplementary Figure 6.

**a and b** shows the XYZ projection of the dead *M. smegmatis*(stained with DAPI) mediated changes.

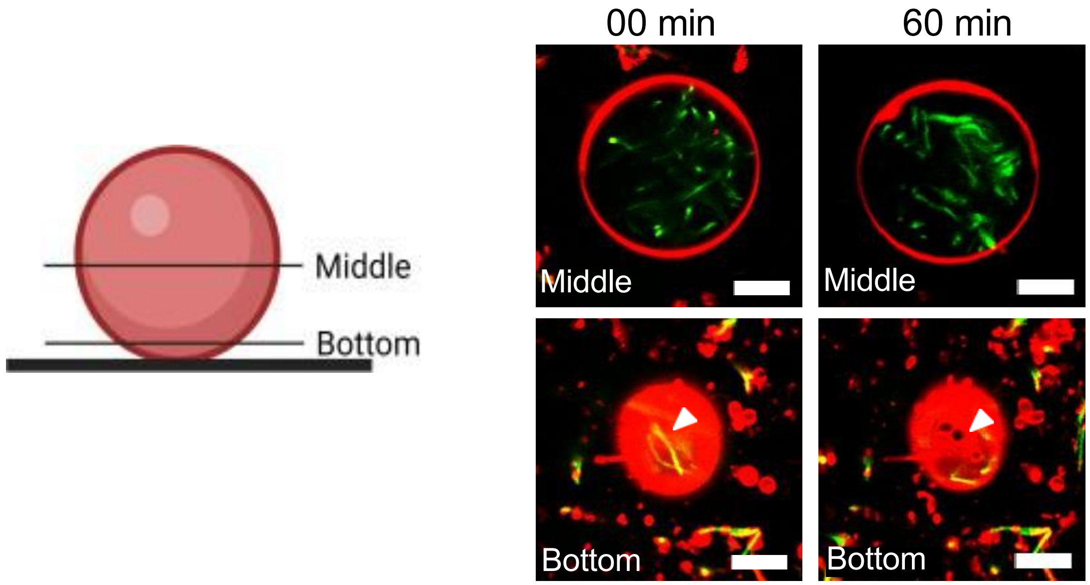


Supplementary Figure 7.
Schematic representation and XY plane of the same GUV from main Fig.2d, representing different planes.

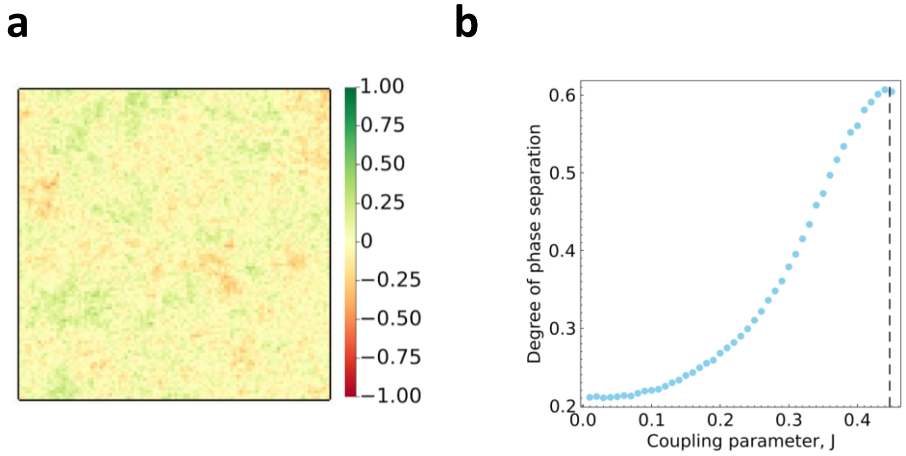


Supplementary Figure 8.

**a.** Time-averaged state of the membrane. Lipids do not show phase separation in the absence of any bacterial contacts ($N_c$ = 0). **b.** The degree of phase separation increases as the coupling parameter, *J* approaches the demixing critical point (shown by the dashed line). (*p*=2, $N_c$=300).

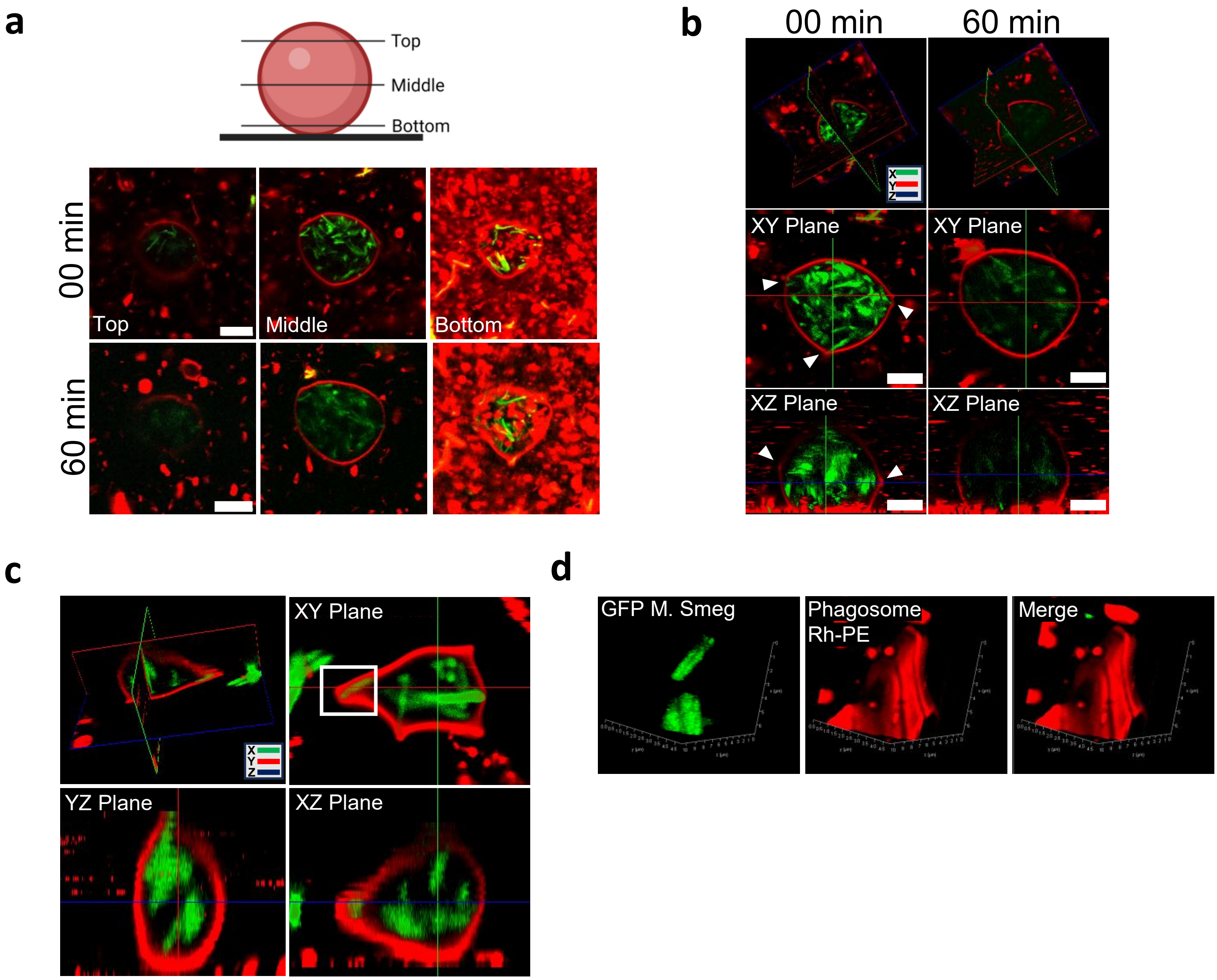


Supplementary Figure 9.

**a.** Schematic representation and XY plane of the same GUV from main Fig 3b, representing different planes. **b.** XYZ Projection of the mycobacteria load mediated changes from main Fig 3b representing the budding event over time. **c.** XYZ Projection of the mycobacteria load mediated changes representing the deformation in the membrane. Scale bar, 10µm. **d.** 3D Zoomed image of confocal images of region near to direct contact of bacteria from the c.

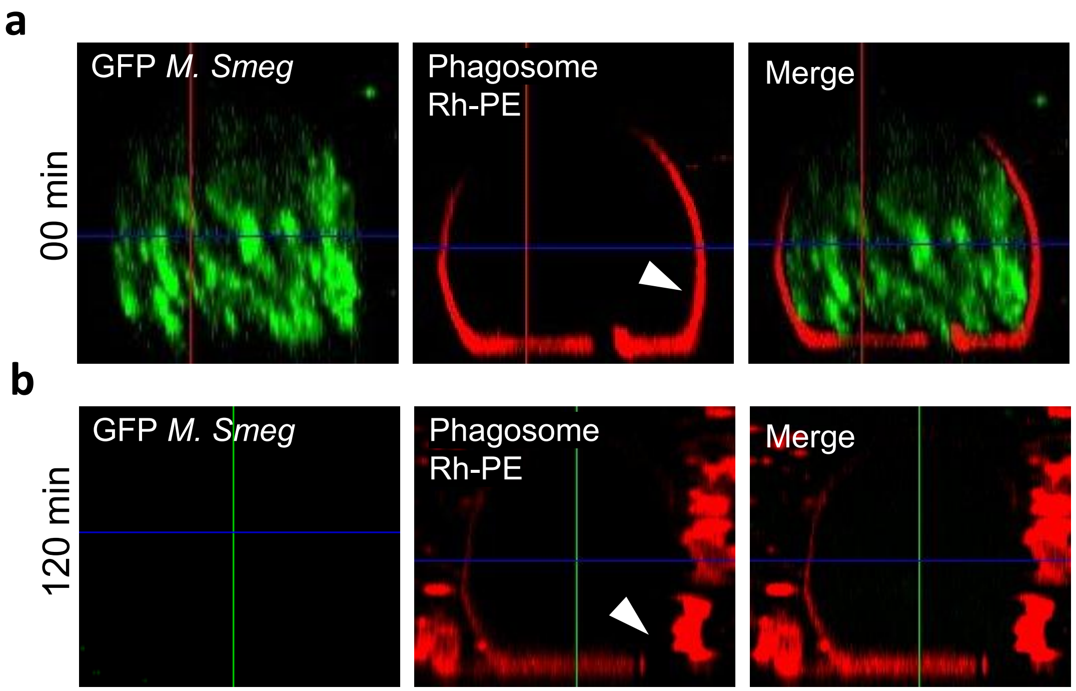


Supplementary Figure 10.
XYZ projection of higher load of bacteria as shown in main Fig 4i at before (**a**) and after (**b**) addition of ESAT-6 respectively.
