## Supplementary text for "Mycobacterial load and direct contact primes spacious to compact phagosome switching for egress"

### Supplementary Movie Legends

Movie. S1.
3D-reconstruction of confocal images of Mtb-mCherry (red) inside Lamp1-positive vesicles (green) inside THP1 cells. The images were acquired in the Olympus FV3000 system and reconstructed using Imaris Viewer 10.2.0 software where the rotation is initiated after zooming into the entrapped bacteria at a scale of around 4-6um. Compact vesicle formation can be seen around the bacteria where there is close association of LAMP1 with the Mtb both in the video as well as the representative snapshots.

Movie. S2.
3D-reconstruction of confocal images of Mtb-mCherry (red) inside Lamp1-positive vesicles (green) inside THP1 cells. The images were acquired in the Olympus FV3000 system and reconstructed using Imaris Viewer 10.2.0 software where the rotation is initiated after zooming into the entrapped bacteria at a scale of around 4-6um. Spacious vesicle formation can be seen around the bacteria where there is a comparatively loose association of LAMP1 with the Mtb as there is a pocket for the bacteria but the volume of the vesicle seems to exceed that of the Mtb which emphasized by the excess space inside the vesicle.

Movie. S3.
Time-lapse confocal imaging of phagosomal membrane mimic GUVs labelled with 0.1% Rhodamine-PE (Rh-PE), encapsulating live bacteria, were immobilized using 0.01% Biotinylated-PEG. Upon monitoring over 60 min (0.0508 fps). Scale bar, 10 µm. with accelerated Video.

Movie. S4.
Time-lapse confocal imaging of GUVs of phagosomal membrane composition (without bacteria) labelled with 0.05% Rh-PE (red) and 0.05% Top Fluor cholesterol (green) and immobilized using 0.03% PEG-Biotin. Upon monitoring over 60 min (0.0083fps). Scale bar, 10 µm. with accelerated Video.

Movie. S5.
Time-lapse confocal imaging of GUVs of phagosomal membrane composition labelled with 0.05% Rh-PE (red) and 0.05% Top Fluor cholesterol (green) and immobilized using 0.03% PEG-Biotin encapsulated with live M. Smegmatis (green). Upon monitoring over 60 min (0.0083fps). Scale bar, 10 µm. with accelerated Video.

Movie. S6.
Time- lapse confocal imaging of GUVs of phagosomal membrane composition labelled with 0.1% Rh-PE (red) and immobilized using 0.03% PEG-Biotin mimicking spacious phagosome encapsulated with live *M. smegmatis* (green). Upon monitoring over 60 min (0.0083fps). Scale bar, 10 µm. with accelerated Video.

Movie. S7.
Time- lapse confocal imaging of GUVs of phagosomal membrane composition labelled with 0.1% Rh-PE (red) and immobilized using 0.03% PEG-Biotin mimicking Compact phagosome encapsulated with live *M. smegmatis* (green). Upon monitoring over 60 min (0.0083fps). Scale bar, 10 µm. with accelerated Video.

Movie. S8.
Time- lapse Epi-fluorescence imaging of GUVs of phagosomal membrane composition labelled with 0.1% Rh-PE (red). Scale bar, 10µm. Aspirated with Pressure (Δp=50Pa) and monitored over the time.

Movie. S9.
Time- lapse Epi-fluorescence imaging of GUVs of phagosomal membrane composition labelled with 0.1% Rh-PE (red) mimicking spacious phagosome encapsulated with (green). Scale bar, 10µm. Aspirated with Pressure (Δp=50Pa) and monitored over the time.

Movie. S10.
Time- lapse Epi-fluorescence imaging of GUVs of phagosomal membrane composition labelled with 0.1% Rh-PE (red) mimicking Compact phagosome encapsulated with live *M. smegmatis* (green). Scale bar, 10µm. Aspirated with Pressure (Δp=50Pa) and monitored over the time.

Movie. S11.
Time- lapse confocal imaging of GUVs of phagosomal membrane composition labelled with 0.1% Rh-PE (red) and immobilized using 0.03% PEG-Biotin incubated with 2.5µM doped FITC-ESAT-6 (green). Upon monitoring over 120 min (0.0667 fps). Scale bar, 10 µm. with accelerated Video.

Movie. S12.
Time- lapse confocal imaging of GUVs of phagosomal membrane composition labelled with 0.1% Rh-PE (red) and immobilized using 0.03% PEG-Biotin mimicking spacious phagosome encapsulated with live *M. smegmatis* (green) and incubated with 2.5µM doped FITC-ESAT-6 (green). Upon monitoring over 120 min (0.0667 fps). Scale bar, 10 µm. with accelerated Video.

Movie. S13.
Time- lapse confocal imaging of GUVs of phagosomal membrane composition labelled with 0.1% Rh-PE (red) and immobilized using 0.03% PEG-Biotin mimicking Compact phagosome encapsulated with live *M. smegmatis* (green) and incubated with 2.5µM doped FITC-ESAT-6 (green). Upon monitoring over 120 min (0.033 fps). Scale bar, 10 µm. with accelerated Video.
