## Supplementary text for "Mycobacterial load and direct contact primes spacious to compact phagosome switching for egress"

**Model for bacterial contact-induced phase separation in a membrane**

We model the membrane using a two-dimensional Ising model on a square lattice of size NxN, with periodic boundaries. The two spins represent the liquid-ordered and liquid-disordered phases of the membrane. The spins interact with their nearest neighbour via the coupling parameter J. To account for bacterial contacts on the membrane, we introduce patches of size *p*x*p*, where the membrane has a strong likelihood of being in the liquid-disordered state. This is imposed through the local field *h*. The Hamiltonian for the system is

$$H = -J \sum_{\langle i,j \rangle} s_i s_j - h \sum_{i \in P} s_i$$

, where the angled brackets represent the nearest neighbour pairs and *P* represents the patches.

The interaction parameter *J* and local field *h* are in units of $k_BT$. We carry out Monte-Carlo simulations following conserved order-parameter (nonlocal Kawasaki) dynamics. For each parameter set, the system is evolved for $10^4$ to $10^5$ Monte Carlo sweeps. The patches representing bacterial contacts are placed randomly on one half of the membrane. This is done since it is observed that a significant amount of Rh-PE label moves to the bottom of the GUV, where presumably most of the bacteria make contact. The degree of phase separation, for a given parameter set, is calculated as the sum of absolute values of the time-averaged state normalized by the total number of spins. A value close to 0 implies that the system is well-mixed and a value close to 1 means that the system is strongly phase separated.

| **Parameter used** | **Range/Value** |
|---|---|
| Lattice size, N | 100 |
| Fraction of up spins, ϕ | 0.5 |
| Local field, $h/k_BT$ | 5.0 |
| Coupling parameter, $J/k_BT$ | 0.01-0.45 |
| Contact patch size, *p* | 2-20 |
| Number of contact patches, $N_c$ | 0-2000 |
